# Parallel lemniscal and non-lemniscal sources control auditory responses in the orbitofrontal cortex (OFC)

**DOI:** 10.1101/2020.02.05.935569

**Authors:** Hemant K Srivastava, Sharba Bandyopadhyay

**Affiliations:** Advanced Technology Development Centre; Department of Electronics and Electrical Communication Engineering, Indian Institute of Technology Kharagpur, Kharapur, India 721302

**Keywords:** deviant detection, persistent activity, stimulus history dependence, lemniscal non-lemniscal auditory pathways, stimulus outcome association

## Abstract

The orbitofrontal cortex (OFC), controls flexible behavior through stimulus value updating based on stimulus outcome associations, allowing seamless navigation in dynamic sensory environments with changing contingencies. Sensory cue driven responses, primarily studied through behavior, exist in the OFC. However, OFC neurons’ sensory response properties, particularly auditory, are unknown, in the mouse, a genetically tractable animal. We show that mouse OFC single neurons have unique auditory response properties showing pure deviance detection and long timescales of adaptation resulting in stimulus-history dependence. Further, we show that OFC auditory responses are shaped by two parallel sources in the auditory thalamus, lemniscal and non-lemniscal. The latter underlies a large component of the observed deviance detection and additionally controls persistent activity in the OFC through the amygdala. The deviant selectivity can serve as a signal for important changes in the auditory environment. Such signals if coupled with persistent activity, obtained by disinhibitory control from the non-lemniscal auditory thalamus or the amygdala, will allow for associations with a delayed outcome related signal, like reward prediction error, and potentially forms the basis of updating stimulus outcome associations in the OFC. Thus the baseline sensory responses allow the behavioral requirement based response modification through relevant inputs from other structures related to reward, punishment, or memory. Thus, alterations in these responses in neurological disorders can lead to behavioral deficits.

## Introduction

The OFC, a part of the prefrontal cortex (PFC), is involved in flexible behaviour (Miller and Cohen, 2001; Wallis, 2007; Fritz et al., 2010) by encoding specific stimulus–outcome or action-outcome expectancies as well as by dynamically revaluing such expectancies on the basis of behavioural demands and motivational states (Delamater, 2007; Rudebeck et al., 2008; Wilson et al., 2014; Fresno et al., 2019). Specific OFC circuits can control specific aspects of flexible behaviour and multiple reinforcement learning processes (Lee et al., 2012; Groman et al., 2019). In order for OFC neurons to encode the sensory attributes and subjective value of outcomes associated with external stimuli (Schoenbaum and Roesch, 2005; Delamater, 2007; Ostlund and Balleine, 2007) it requires sensory inputs. It is known that sensory stimulus-evoked signals in the OFC can distinguish between appetitive and aversive outcomes (Morrison and Salzman, 2011) associated with the stimuli. Further, the OFC also has the capacity to influence sensory processing by modulating neuronal receptive fields in early sensory cortices, particularly the auditory cortex (Winkowski et al., 2013).

In order to understand how mechanistically specific stimulus outcome associations are created and how the stimulus evoked OFC responses may influence sensory representation, it becomes crucial to delve into the origins of sensory inputs and sensory response properties of the OFC. In the case of auditory stimuli, the pathways involved and their contribution to auditory responses in the OFC are not known. What aspects of information in the ongoing auditory environment, how and in what form reaches the OFC would determine how mechanistically stimulus-outcome expectancies or values would get computed or updated. As a first step, we consider the auditory evoked responses of OFC neurons, purely from a sensory perspective and attempt to decipher the crucial components of the auditory pathway involved in shaping auditory responses and their properties in the OFC.

With single unit recordings in both awake and anaesthetized OFC we show that auditory responses in the OFC are strongly context dependent with long timescale history dependence and pure deviance detection, unlike the auditory pathway. Investigation of anatomical and functional sources of inputs show that both the lemniscal and non-lemniscal pathways at the cortical and sub-cortical levels shape auditory responses in the OFC. With multiple pharmacological inactivation experiments, the contributions of multiple auditory cortical and subcortical areas in OFC’s auditory responses were assessed. In the auditory cortex (AC), the dorsal region (AuD), with the most projections to the OFC, surprisingly did not contribute to auditory responses of the OFC, while the other higher order non-lemniscal ventral auditory area, AuV (Sacco and Sacchetti, 2010) was found to be the main source of auditory evoked excitatory drive to the OFC. The primary auditory cortex (A1) contributed to temporal response properties in the OFC. Further, considering the auditory thalamic (medial geniculate body, MGB) sources showed that non-lemniscal AuV’s contribution to OFC responses originate from the lemniscal MGBv through its direct projections to AuV (Ohga et al., 2018). The non-lemniscal, polymodal medial division of MGB, MGBm (Weinberger, 2011; Lee, 2015) inactivation however caused the OFC auditory responses to become persistent, like frontal cortex responses during working memory dependent tasks (Fuster and Alexander, 1971; Funahashi et al., 1989; Schoenbaum and Setlow, 2001). Thus MGBm could be an important component of the circuit that allows responses to persist for a longer duration depending upon task demands. Thus the MGBm is a source that causes auditory driven long lasting inhibition in the OFC. Inactivation of BLA, providing inhibitory inputs (Dilgen et al., 2013; McGarry and Carter, 2016; Lichtenberg et al., 2017) to the OFC and known to receive MGBm inputs via the LA, lateral amygdala (Ledoux, 2000; Woodson et al., 2000), showed similar emergence of auditory driven persistent activity. Thus the MGBm-LA-BLA-OFC pathway provides crucial inhibitory control on AuV driven auditory responses in the OFC. Further, the same pathway contributed substantially to the strength of deviant selectivity in the OFC. We suggest that the feedforward inhibition (Dilgen et al., 2013) from BLA to OFC and parallel MGBm to LA inhibition (Woodson et al., 2000) transmitted to OFC via BLA allow for two independent controls to generate persistent activity in the OFC required for stimulus outcome associations.

## Methods

### Animals

All animal experiments were approved by Institutional Animal Ethics Committee (IAEC) of Indian Institute of Technology Kharagpur. Animals were reared under a 12-h light/dark cycle and maintained at a temperature of 22–25 °C and had access to food and water *ad libitum*. C57BL/6 mice of either sex aged between P-25 to P-45 were used for the experiments. Data acquired in pharmacological block experiments before blocking were also included in analysis of OFC responses.

**Table 1.** Experiment wise animals used

| Experiment | Number of mice |
| --- | --- |
| OFC Electrophysiology | 27 |
| AC Electrophysiology | 5 |
| Anatomy | 9 |
| A1 Inactivation (+ Bilateral) | 7 (+ 2) |
| AuV Inactivation | 5 |
| AuD Inactivation | 3 |
| MGBv Inactivation | 2 |
| MGBm Inactivation | 5 |
| BLA Inactivation | 5 |
| Awake OFC Electrophysiology | 5 |
| **Total** | **74** |

### Animal Preparation

#### Anesthetized recordings

Animals were anesthetized using Isoflurane (5% for induction and around 1-1.5 % for maintenance). Body temperature was maintained at 39 °C by placing the animal on a heating plate. A small incision was made to expose the skull and a metal plate was attached on to the skull to head fix the animal. Once head fixed, a small (∼2 mm dia) craniotomy was performed to remove the skull over the recording site. For OFC, the stereotaxic coordinates used were AP=+2.5 mm ML=1 mm from the Bregma, DV=1.8mm (Paxinos and Franklin, 2013) from the brain surface. For AC, the recording site was identified based on vasculature (Sawatari et al., 2011). All recordings were performed with a microelectrode array (MEA) with a 4X4 grid (125 μm between rows and columns) of epoxy-coated tungsten electrodes (MicroProbes, impedance ∼3-5 mega ohms).

#### Awake head-fixed recordings

As in the anesthetized case, a similar but smaller craniotomy (∼1 mm dia) was performed and the electrodes were advanced into the recording site and the held fixed on the skull using Metabond (C & B superbond). A titanium plate was also fixed to the skull (posterior to the electrodes) with Metabond to head-fix the animal during the experiments. The Animals were allowed to recover for 5 days and then were habituated with the recording setup for 30 min for three days before data collection. Data collection lasted for less than 7 days, with ∼1 hour long sessions every day. Units collected on each day from the recording electrodes were considered as separate units.

#### Stimulus

All acoustic stimulation was presented from the right side, contralateral to the recording site. Initially noise bursts (6-48 kHz bandwidth, 50 ms, 5 second gaps, of multiple intensities – 40 dB to 0 dB attenuation, in 10 dB steps; 0 dB attenuation corresponds to ∼95 dB SPL for tones) were used to obtain threshold sound level for noise. Next single units were characterized with pure tones (50 ms, 6-48 KHz, 1/2 octave apart, 70-80 dB SPL – depending on noise threshold, 5 seconds apart, except mentioned otherwise) to obtain tuning and the best frequencies (BF) at the chosen sound level. Next response to a pair of oddball stimulus set was collected. The oddball stimulus consisted of a standard token (S, 50 ms, either a noise token or a pure tone) and deviant token (D, 50 ms, either a pure tone or a noise token respectively for noise-tone, NT or TN, oddball; S and D were both pure tones in case of tone-tone, TT, oddball). The S-D stream had 15 tokens presented usually at 4 Hz or 3.3 Hz; all the tokens were S tokens except the 8^th^ token (usually) which was the D token. In the second of the pair of oddball stimulus set the S and D tokens were swapped. Each oddball set was repeated 20-30 times with a gap of >5s between each repetition. All sound tokens presented in all kinds of stimuli had 5ms rise and fall times. Since recordings were with MEAs with 16 electrodes, the pure tone frequency was chosen based on the tuning of the majority of simultaneously recorded neurons, such that the chosen tone frequency was within the pure tone tuning range of most neurons. The stimuli were generated using a custom written software in MATLAB (Mathworks) and presented with Tucker Davis Technologies (TDT) ES1 speakers (driven with TDT ED1 drivers) after generation with a TDT RX6 processor and attenuated using a TDT PA5. The speaker was placed 10 cm away from the contralateral ear.

### Electrophysiology

#### Anesthetized

The MEA was slowly advanced in to the recording site with the help of a manipulator (MP-225, Sutter). The electrodes were allowed to settle and stabilize for about 30 min. before the data was acquired. Data were collected using custom written software (MATLAB), through a unity gain headstage (16 channels, Plexon HST 16o25) amplifier, followed by a preamplifer (PB3, Plexon, 1000X). Wide-band neural signals (0.7–8 kHz) as well as a parallel set of 16 channels with spike signals (150 Hz to 8 KHz) were stored after digitizing at 20 kHz using a A/D board (National Instruments). Off-line analysis was performed with stored data. At the end of the experiment, the animal’s brain was harvested for *post hoc* examination of the recording site.

#### Awake

For awake recordings, the animal was placed in a small tube with head protruding out and fixed using the titanium plate implanted during surgery. Rest of the procedure was similar to the anesthetized recordings.

#### Anatomy

9 animals were injected with 200 nl of green retrobeads (Lumafluor) into OFC and 3 of them were also injected with 100 nl of anterograde tracing AAV.CB7.CI.mCherry in MGBv using Nanoject II. After 14 days of injection, the animals were transcardially perfused with 20 ml PBS followed by 20 ml of 4% Paraformaldehyde and brain was harvested and kept in 4% paraformaldehyde overnight. 100 µm thick brain sections were cut using a vibratome (Leica VT1000S), mounted on a glass slide with fluomount cover slip, and observed under a fluorescence microscope (Leica DM2500).

#### Electrophysiology with pharmacological inactivation

A small burr hole was made on the skull over the area to be inactivated, ipsilateral (or both sides as mentioned) to the recording site. A Hamilton syringe (7000 series) loaded with the 200nl of GABA agonists (5μg/μl muscimol and 2μg/μl baclofen) or equal volume saline was inserted and held via a cannula implanted on the skull with dental cement. Only after the syringe was positioned securely, the electrodes were inserted into the recording site as described above. For injecting the agonists/saline during the experiment, the Hamilton syringe was gently pressed/tapped multiple times over 5-10 s period to release the entire volume. There was a wait period of 30 mins for the agonists to have their effect before the next data set was acquired. SR101 was added to the mixture of agonists or saline for *post hoc* confirmation of the target site and spread of the injection. Different divisions of AC were identified and marked based on the vasculature.

For MGB (Slater et al., 2019) and BLA (Luna and Morozov, 2012) following stereotaxic coordinates were used: MGB: AP= −3.27 mm, ML= 2.0 mm from Bregma, −3.0 mm from the brain surface BLA: AP= −1.3 mm, ML=3.2 mm from Bregma, 3.8 from the brain surface.

### Data Analysis

Spike Sorting was done offline in custom written MATLAB software. Data were notch filtered (Butterworth 4^th^ order) to reject any remnant power supply 50Hz oscillations. Spiking activity was obtained directly from the spike channels of the PBX3 preamp. Waveform fluctuations above 3.5-4 standard deviations (usually 4) from the baseline were isolated and based on shapes, spike waveforms were clustered into different groups. The timing of spikes with respect to data collection onset (and hence also stimulus presentation) were extracted for each spike shape (single unit) for further analysis. A single unit was considered as responsive if the spike rate within 400 ms (200 ms in case of awake condition) of stimulus presentation was significantly different from the baseline (300 ms preceding the stimulus, t-test, alpha=0.05). Response latency was calculated as the time at which the spike rate in the average peristimulus time histogram (PSTH, 20ms bins) was maximum.

#### Narrow tuning and calculation of BF

Tuning of neurons was considered to be narrow (well-tuned) or bimodal and/or broad. The frequency corresponding to the maximum response rate out of the 7 frequencies presented to narrowly-tuned units (below) was considered the BF of the unit. Those units were narrow-tuned whose average response to frequencies other than one octave around BF was two standard deviations (variability in response rate at BF) below the response at BF.

#### Spatial BF Variability

The standard deviation of BFs of all the simultaneously acquired units (only narrowly tuned, above) was normalized by the product of area accommodating these responding units and the number of units.

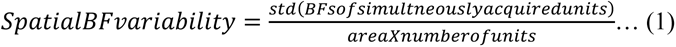

For simulating a distribution of completely heterogeneous BFs, each unit was randomly assigned a BF (uniformly over the 7 frequencies used) and BF variability was calculated with 1000 bootstraps.

#### CSI calculation

For common selectivity index (CSI) calculation, those units were included which responded to at least one of the four stimulus tokens, first of each of the S tokens in the oddball pair S_X_ and S_XS_ (XS being the swap of the X oddball), and the deviant tokens, D_X_ and D_XS_. CSI was calculated as per the following equation (Ulanovsky et al., 2003)

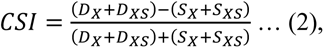

where, *S_i_*/*D_X_*_/*XS*_ represent the mean rate response to those tokens – *S_i_* the *i^th^* token and *S_PT_* being the token preceding the *D* token. The rate responses in each case were computed based on the following windows:

-For S_1_ (100-400 ms from S_1_)

-For S_ALL_ (S_2_ +100 ms - till deviant; entire length)

-For S_PT_ (100ms from S_PT_ - up to deviant)

-For D (100-400 ms from deviant)

When a sufficient sized population (at least 50 units) of paired data was not available (as in the case of tone-tone, TT oddball and in before versus after pharmacological inactivation experiments) deviant selectivity index (DSI) of each unit was calculated as follows

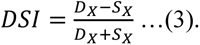

#### Anatomy

The number of retrobeads was quantified using a threshold that was manually set for individual sections depending upon the background intensity. The laminar demarcation was based on distance from pia, which was also corroborated in a subset with the MGBv projections observed with AAV.CB7.CI.mCherry. All beads within 300 µm from the pia were marked as layer 2-3, beads between 300 to 450 µm (or mCherry) were marked as layer 4 and beads below that were grouped into layer 5-6.

Demarcation of AuV, A1 and AuD to confirm correct injections through spread of SR101 in blocking solution was through brain atlas (Paxinos and Franklin, 2013) and distance from Rf (rhinal fissure). Only those animals were included in the dataset whose spread was *post hoc* confirmed as above. Injections in MGBv and MGBm were targeted and *Posthoc* conformed through brain atlas (Paxinos and Franklin, 2013)

#### Response duration

A sliding response window of size 100 ms starting from the stimulus start, in 20 ms steps, was compared with the random 100 ms window in the baseline for significance. Consecutive significant bins with a time difference of less than 100 ms between them were joined together for the determination of response duration.

#### Spike timing jitter

The spike timing jitter expressed as variability in the timing of individual spikes across trials was computed as the reliability (R_corr_) in spike timing which is a measure of similarity between pairs of individual spike trains (Schreiber et al., 2003). The trial wise individual spike trains binned at 1 ms was convolved with a Gaussian filter of σ=2 ms and then coefficient of correlation was calculated between all pairs of trials using the MATLAB function *corrcoef*. Higher the R_corr_ value less is the spike timing jitter.

#### Pairwise correlations

Pairwise correlations were calculated as correlation coefficient using the MATLAB function *corrcoef* between mean PSTHs (from stimulus start) of simultaneously recorded units.

## Results

### OFC neurons respond to sound with very long timescale dependence

OFC single neurons, mainly studied in primates, are known to respond to sounds and other sensory stimuli. However, response properties of particularly auditory stimuli in the mouse OFC are not known. Although mouse OFC single neurons have been shown to respond to a variety of sounds (Winkowski et al., 2017), it is unclear how selective the responses are and how the responses change under different sensory contexts. Most studies have considered auditory cue associated responses in the OFC in the context of behavior with single or multiple cues linked to aspects of behavior. However, in a dynamic natural environment, in which an animal is required to involve the OFC for the purposes of flexible stimulus value guided behavior (Wallis, 2007; Groman et al., 2019), the sensory response properties in different sensory contexts is crucial. To first obtain OFC single neuron responses to sounds, independent of behavioral state, active memory and other cognitive processes, we considered single units in the OFC of the anesthetized mice.

OFC single unit recordings were performed in the anesthetized mouse in response to auditory stimuli. The recording sites were confirmed to be in the OFC (lateral and/or ventral), *post-hoc* by Nissl staining of coronal sections (Fig. 1A). We found that neurons in the mouse OFC robustly responded to sound stimulation (Fig. 1B and 1C) with broadband noise and pure tones (also see Winkowski et al., 2017), with typically monotonic rate intensity functions with noise (Fig. 1B, right) and variety of BFs (Fig. 1Ci-iv) with narrow (Methods, 45% of the responding units) and bimodal to broad tuning (55% of the responding units, Fig 1Cv-vi). While most of the responding units increased their firing rate at BF (∼86%, 816/949 units), about ∼14% (133/949) of units reduced their firing rate at some frequency (Fig. 1D); 76 of the 133 units were excited at BF while the rest showed no excitatory responses. Notably the inhibition observed in all the 133 units (Fig. 1D), were all at either 6 or 8.5 kHz. The mean peak latency observed at the BF of units with excitatory responses was 284 ± 3 ms (Fig. 1e). The latency was substantially longer than what is observed in the mouse AC single unit responses but has similar latencies as a late component of auditory responses (Chen et al., 2015), likely in the nonprimary AC.

**Figure 1.**
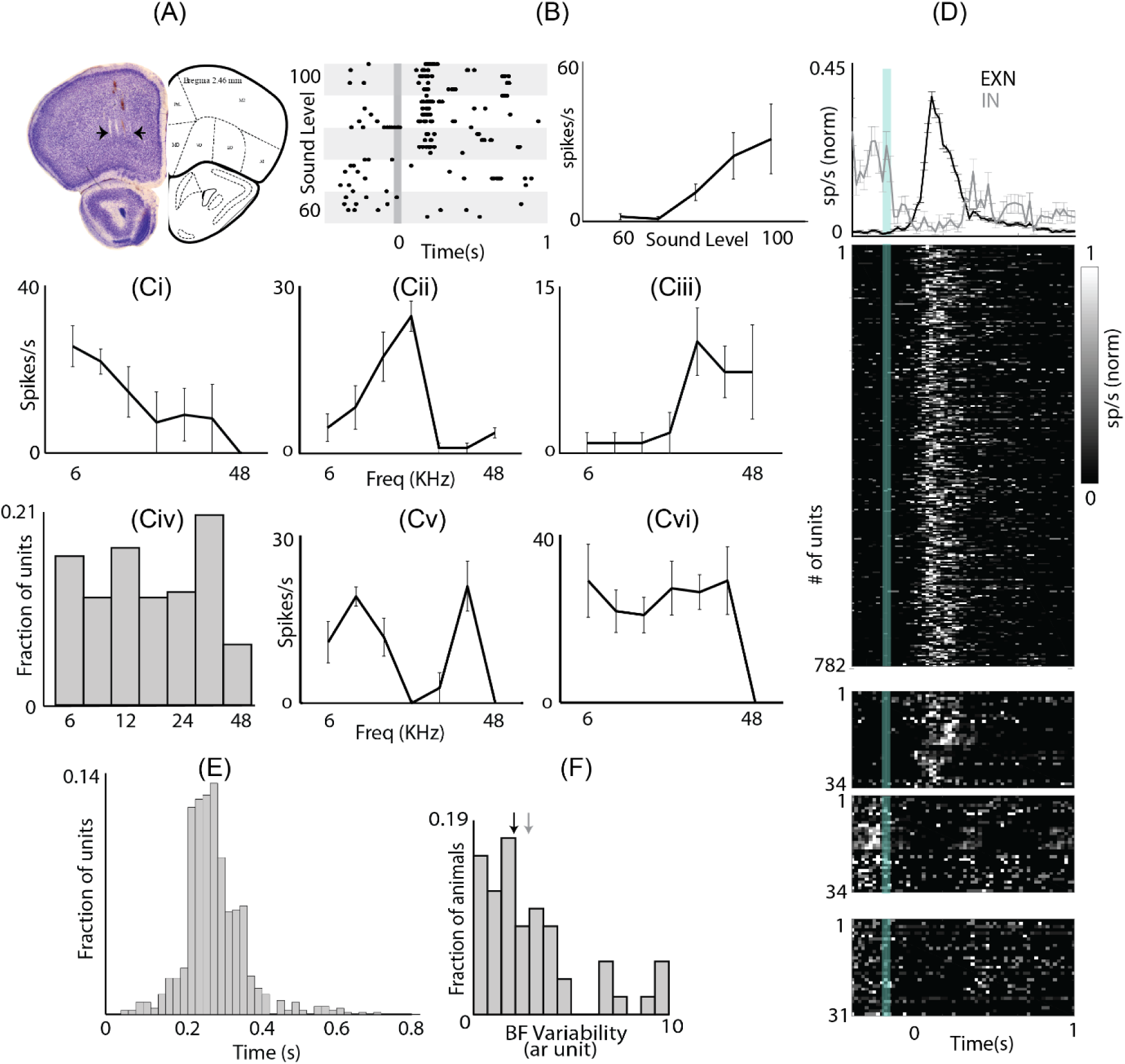
– OFC responds to auditory stimulation. (A) Nissl stain of OFC section showing electrode tracks. Black arrows mark the medio-lateral extent of the electrode array. (B) an example unit showing responses to increasing intensities of broadband noise in raster plot (left) and rate-level curve (right). (Ci-iii) Tuning curves of three example units tuned to low (Ci), mid (Cii) and high (Ciii) frequencies. (Civ) Distribution of units tuned to different frequencies. (Cv) An example unit with bimodal tuning curve. (Cvi) An example unit with broad tuning curve (D) top: mean normalized PSTH ± SEM of all units showing excitation (black) and inhibition (grey) upon auditory stimulation. Bottom; individual unit’s PSTH (upper panel: units showing excitation, middle two panels: units showing excitation to some frequency and inhibition to some other frequency, lower panel: units showing inhibition). (E) distribution of peak response latency to the BF. (F) Spatial BF variability. Black arrow marks the median of the distribution and grey arrow marks the median of the distribution of completely random BF organization.

In order to check for possible topographical organization of tuning based on the narrowly tuned units in the horizontal plane of the OFC, we considered the variability of BF of units recorded simultaneously. The distribution of calculated spatial BF variability in simultaneously recorded units (Eqn. 1, Methods) across all recording locations was compared with the distribution expected with completely random BFs at each recording site (Methods) (Fig 1E, black arrow median of data, gray arrow median of the distribution with spatially random BFs). The two distributions were not significantly different (KS-test, p=0.06). Thus we conclude that the local organization is completely random indicating absence of any BF based organization at the spatial scales (∼400 μm) of our recording along the horizontal plane in OFC.

We used a 5s inter stimulus gap to characterize single unit responses to tones and noise in the mouse OFC. While, stimulus presentation rates in the auditory pathway, to rule out any effects of adaptation, have been long established (Antunes et al., 2010), corresponding repetition rates in the OFC are not known. To investigate effects of adaptation and stimulus history, we recorded OFC responses to pure tone presentation with varying inter-trial interval (ITI) with short interval having less than 5s, mid interval ranging from 5s to 7s and long interval ranging from more than 7s to up to 11s (Fig. 2A and C). Since multiple time scales of adaptation exist (Ulanovsky et al., 2004) in the auditory cortex itself, we investigated the timescales of adaptation in the OFC in comparison to AC (Fig. 2B and F, both A1 and AuV; determined from *post hoc* Nissl stains with electrode tracks). We found that the response profiles of these three ITIs in the OFC were significantly different (one-way ANOVA Fig. 2C&D). The peak spike rate of the long ITI group was significantly different (one-way ANOVA, p<0.001) from the mid and short ITIs (Fig. 2D), whereas the latency (Fig. 2E) of the short group was significantly different from the other two groups (one-way ANOVA p<0.01 between short and long and p<0.001 between short and mid). These results indicate that OFC neurons show very long (at least up to tens of seconds) timescales of stimulus history dependence, reflected in the spike rate or latency of the response. Similar analyses of the AC neurons (Fig. 2F-H) show that neither the peak spike rates (Fig. 2F&G) nor the response latencies (Fig. 2H) were different in the three groups. Thus OFC responses to auditory stimuli were found to have unprecedented temporal stimulus history dependence. Such remarkable dependence of sensory responses on long stimulus history, unlike in the sensory cortex, would be crucial in normal environments with continuously varying sensory inputs, unlike experimental controlled environments.

**Figure 2.**
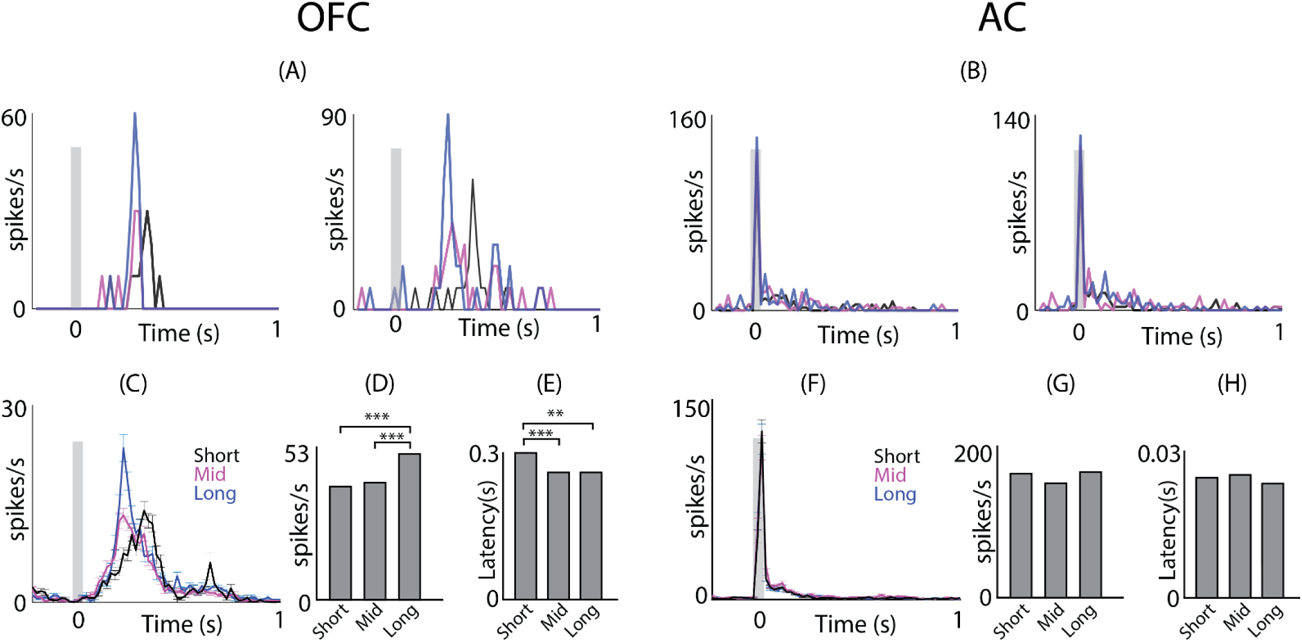
– OFC neurons show long timescales of adaptation (A) Psth of two example units in the OFC at different ITIs (black; less than 5 secs (short), magenta; between 5 and 7 secs (mid), blue; more than 7 secs up to 11 secs (long)). (B) Similar to (A) in the AC (C) mean population PSTH ± SEM at different ITIs in the OFC (D) Mean peak spike rate in the OFC (E) mean peak response latency in the OFC (F) mean population PSTH ± SEM at different ITIs in the AC (G) mean peak spike rate in the AC (H) mean peak response latency in the AC

### Neurons in the OFC show high context dependence and pure deviance detection unlike in the AC

Given the long temporal dependencies, presumably due to strong and long lasting adaptation, it becomes important to find, to what aspects in streams of sounds, neurons in the OFC respond. Since most neurons in the OFC are broadly tuned and respond to noise (Fig. 1), we first considered an oddball stimulus set with noise tokens as the standard (S) stimulus with a tone embedded in the stream as the D token (Methods). Thus responses of OFC single units to a pair of NT and TN oddball stimuli were collected. We found that OFC neurons robustly responded to the D token (Fig. 3A). Typically, a strong onset response to the first of the standards (a deviant/change from the pre-stimulus silence) was seen, followed by a strong response to the D. Following both the first S and D, the responses adapted quickly without responding to the succeeding tokens. Thus the responses in the OFC to odd-ball stimulation show strong and fast stimulus specific adaptation (Taaseh et al., 2011; Nieto-Diego and Malmierca, 2016).

**Figure 3.**
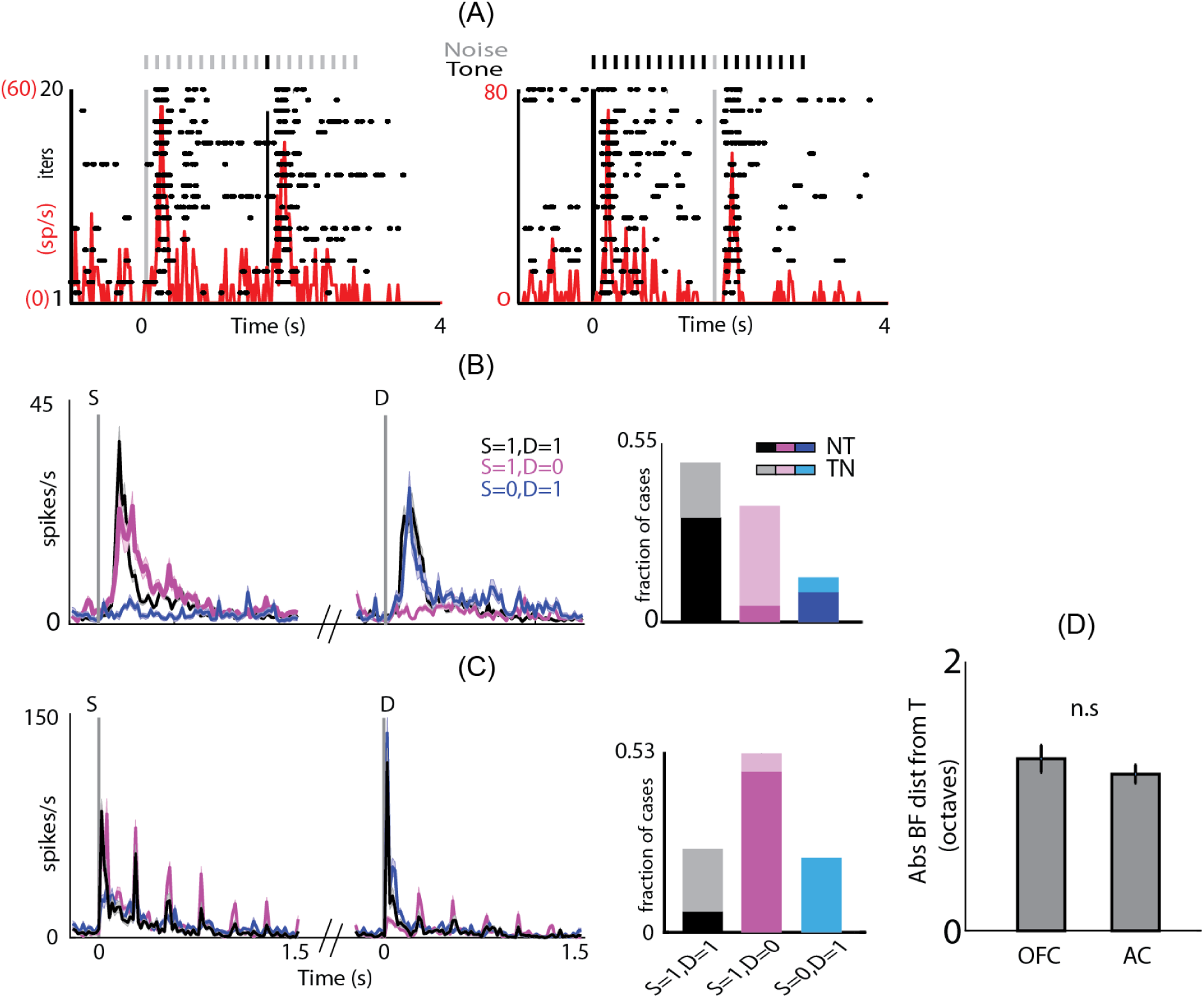
– Pure Deviant Detection in OFC unlike AC (i) (A) An example unit showing auditory response (raster (black) and PSTH (red) to an odd-ball stimulus where noise is played as standard and tone as deviant (NT; left) and its swap (TN; right). The vertical lines mark the onset of standard and deviant (grey for noise and black for tone). (B) (left) Mean population PSTH ± SEM of units responding to both standard onset and deviant (S=1, D=1; black), responding only to standard onset (S=1, D=0; magenta) and responding only to deviant (S=0, D=1; blue). Right: Fraction of units belonging to different categories as described in (B). The darker shades show the fraction of units belonging to NT and lighter shades show fraction of units belonging to TN group. (C) Similar to (B) in the AC. (D) mean (|BF-Tf|) in the NT-TN oddball stimuli in the OFC and AC

Oddball stimulus response pairs (NT and TN) from 109 OFC units (10 animals), 218 cases, where the unit responded at least one of the 4 stimulus tokens (S_1_ and D in each of the two in a pair, Methods). In 36% (79/218) of the cases units responded to both the first S token (S_1_) and the D token (S_1_=1, D=1; 62/79 NT and 17/79 TN cases, Fig. 3B, black), while in 31% (68/218) cases units responded to S_1_ but not D (S_1_=1, D=0; 3/68 NT and 65/68 TN cases Fig. 3B, magenta). Further, 17% (37/218) responded to only the D token (S_1_=0, D=1; 21/37 NT and 16/37 TN cases Fig. 3B, blue) and the remaining 16% did not respond to either. In the last 16% cases there was however a response to one of the stimulus tokens (S_1_ and/or D) in the corresponding swap oddball stimulus. Thus, although OFC neurons were found to be generally responsive to both, the tone token and the noise token (Fig. 1, Methods), their response to the same sound token changed depending on the context of the stimuli. We find that when a neuron responded to a tone as S_1_ (82/109), it was less likely to respond to the noise token as D (65/82 and 17/82, p<0.001, *two proportion z-test*). On the contrary, if a neuron responded to the noise token as S_1_=1 (65/109), it was more likely to also respond to the tone token as D (62/65 and 3/65, p <0.001, *two proportion z-test*). Such context specific selectivity is contrary to expectations as a broadband noise as S is likely to adapt all frequency channels thus masking the response to a tone as D embedded in a sequence of noise tokens. The above suggests that the OFC auditory circuitry is inherently more selective to detecting narrowband sounds in broadband background noise. However, when considering the response of the same unit to a stimulus token (noise or tone) as the S and as D, we find that out of 109 units, 75(17), 7(48), 8(16) and 19(28) units responded to the tone (noise) both as S and D, only as S, only as D and neither respectively. The above suggests that responses to the tone was less context sensitive than noise with more units responding to tones than to noise (75/109 and 17/109, p<0.001, *two proportion z-test*) independent of its occurrence as S or D. This further corroborates the fact that the OFC neurons inherently are more responsive to tones and hence likely narrowband tonal sounds like mouse vocalizations.

In contrast, we found nearly an opposite pattern of context dependence in the AC. Responses to oddball stimuli pairs (NT and TN) from a total of 62 units (51 in A1, 11 in AuV, 124 cases, Fig. 3C) were collected from the AC that showed responses to at least one of the 4 stimulus tokens (as above, Methods). In 22% (28/124) of the cases units responded to both S_1_ and D (7 NT and 21 TN, bar plot Fig. 3C, black), while in 48% (60/124) cases units responded to S_1_ but not D (54 NT 6 TN, bar plot in Fig. 3C, magenta). Further, 20% (25/124) responded to only the D token (none NT and 25 TN, bar plot in Fig. 3C, blue) and the remaining 10% did not respond to either. As expected from adaptation along frequency channels, when there was a response to the noise as S very few units showed a response to the tone as deviant (7/61) and most were unresponsive to the tone (54/61), opposite of what was observed in the OFC. Considering responses of the same neuron to the tone or noise token in either of the two contexts (as S_1_ or D), 7(45), 20(16), 0(1) and 35(0) out of 62 AC units responded to tone (noise) as both S_1_ and D, only as S_1_, only as D and neither of the two. As above, in the AC, contrary to observations in the OFC, the tone responses independent of S and D were absent, while noise responses occurred almost independent of S and D (7/62 and 45/62, p<0.001, *two proportion z-test*). There were many more cases in the AC when there were no responses to the tone either as S or as D. Such units were present in both A1 and AuV in similar proportions (30/51 and 5/11, NS). The choice of tone frequency (Tf) for oddball sets (NT & TN) with respect to the BF of neurons whose responses were collected, could not explain the difference of context dependence in the AC and OFC as in both cases the choices were similar (Fig 3D mean(|BF-Tf|) 1.3 octaves OFC and 1.1 octave AC, NS, *unpaired t-test*).

To find the neuron’s inherent preference to the deviant either T or N, we calculated CSI in three different ways based on the choice of S tokens considered for S_X_ and S_XS_ in Equation 2. We compared the spike rates in response to either the first token of standard (S_1_) and D, or all standard tokens (S_ALL_) preceding D except S_1_ or just the previous token (S_P_) to D. Comparison of responses in scatter plots (Fig. 4A) of spikes rates of S_1_ and D (left), S_ALL_ and D (middle) and S_P_ and D (right) for NT and TN (blue and orange respectively) lack of responses to S beyond S_1_. Further, the different CSI distributions (Fig. 4A, bottom) show that the OFC neurons respond strongly to S_1_ as compared to D probably because S_1_ is a bigger change than D in the stimulus space. The scatter plots of spike rates and CSI indices of S_ALL_ and D and S_P_ and D look very similar suggesting that OFC neurons do not respond to any S except S_1_. We calculated the CSI of OFC neurons by either taking S_ALL_ and D or S_P_ and D, we found that these two CSI distributions were not significantly different, whereas in AC these two distributions were significantly different (p<0.001, *t-test*) (Fig. 4B). This clearly shows a strong adaptation onset right from the second token (beginning of repetition) in OFC unlike ACX suggesting a hierarchy of SSA strength along this pathway.

**Figure 4.**
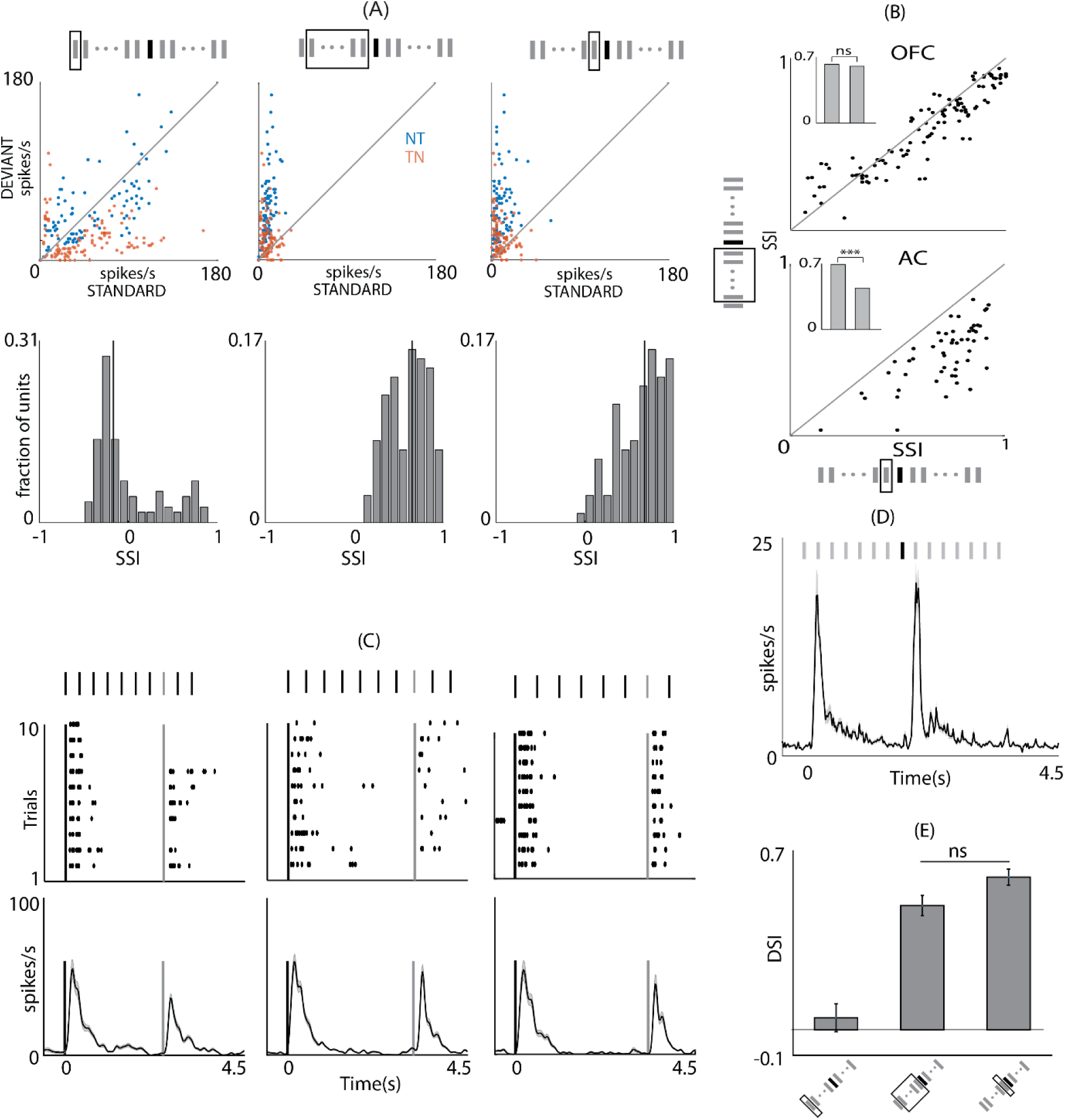
– Pure Deviant Detection in OFC unlike AC (ii) (A) Top: Scatter plot of mean spike rate at standard and deviant in NT (blue) and TN (orange). The spike rates for standard were calculated either considering the S_1_ (left), or S_ALL_ (middle) or S_PT_ (right). The tokens considered in the two cases are enclosed in the grey rectangle. The histograms at the bottom show the CSIs computed by using three different standards as described in (A). (B) Scatter plot of CSIs in OFC (top) and AC (bottom) calculated by taking S_ALL_ and S_PT_. The mean CSIs in the two cases are shown in the bar plots in the inset. (C) Top: Raster plots of example units in response to odd-ball stimulus with different inter-token interval;300 ms (left), 400 ms (middle) and 500 ms (right). Bottom: mean population PSTH for these intervals. (D) Mean population PSTH to tone-tone odd-ball stimulus. (E) Mean DSI for tone-tone odd-ball stimulus with three different standards as described in (A).

Since neurons in the OFC did not respond to any of the S tokens following the response to S_1_, showing pure deviance detection by responding only to the D token (with S_1_ being also effectively a D from the preceding silence), instead of the 200 (usually)/ 250 ms inter token interval (IToI), we also considered IToIs of 300 (27 units), 400 (41 units) and 500 ms (40 units). Single neuron raster plots and population mean PSTHs of responses to oddball stimuli (Fig. 4C) showed no responses to any of the sound tokens other than S_1_ and D, again showing pure deviance detection like responses.

In order to consider the generality of the pure oddball detection with two narrowband sounds instead of one narrowband and one broadband, we also collected responses to oddball stimuli with 2 tones (TT; 102 units) but not necessarily in pairs as in the NT/TN cases. As with TN and NT cases, there was a lack of responses to the S token beyond S_1_ and strong response to the D tone (Fig. 4D). Comparing the mean DSI (Fig. 4E) between the 3 ways of computing selectivity index (considering S_X_ in Eqn. 3, to be response to S_1_, S_ALL_ and S_P_), as in the case of NT/TN CSI, showed no significant difference between DSI_ALL_ and DSI_P_ (0.49 and 0.6, NS, ANOVA). In a small subset of units (*n*=25), for which paired data was available with the same stimulus parameters (repetition rate, position of D) as the TN and NT dataset, CSI_ALL_ and CSI_P_ were also not different (0.23 and 0.26, NS, ANOVA). The comparatively lower CSI value was due to lack of responses to some of the tones as D in this dataset or higher spontaneous rate, which also further shows strong adaptation to the S tone. Thus OFC neurons probed with odd ball stimuli show that they inherently detect changes or violations to the regularity in the stimulus space and could be an important attribute required for flexible value updating in the OFC.

### Sparse responses in the OFC of awake, passively listening mice also show deviance detection

In order to confirm that the observed deviance detection in the OFC is also present in neurons in awake, single unit recordings in passively listening mice were performed with electrodes implanted in the left OFC (Methods). As in the anesthetized case, we found robust responses to auditory stimulation, with 73% (162/219) units showing excitatory responses to pure tones (Fig. 5A&B), which is a much higher proportion of neurons than in an earlier study (Winkowski et al., 2017). The neurons showed expectedly much shorter latency to peak (134.5±2.83 ms SEM, Fig. 5C) than in the anesthetized mice, while a small 14% units showed a very short latency (∼14 ms) and were not included in further analyses as they could be sound evoked movement related. As in the anesthetized mice, in the awake mice also, a small fraction of neurons (4%) showed inhibitory responses to tones.

**Figure 5.**
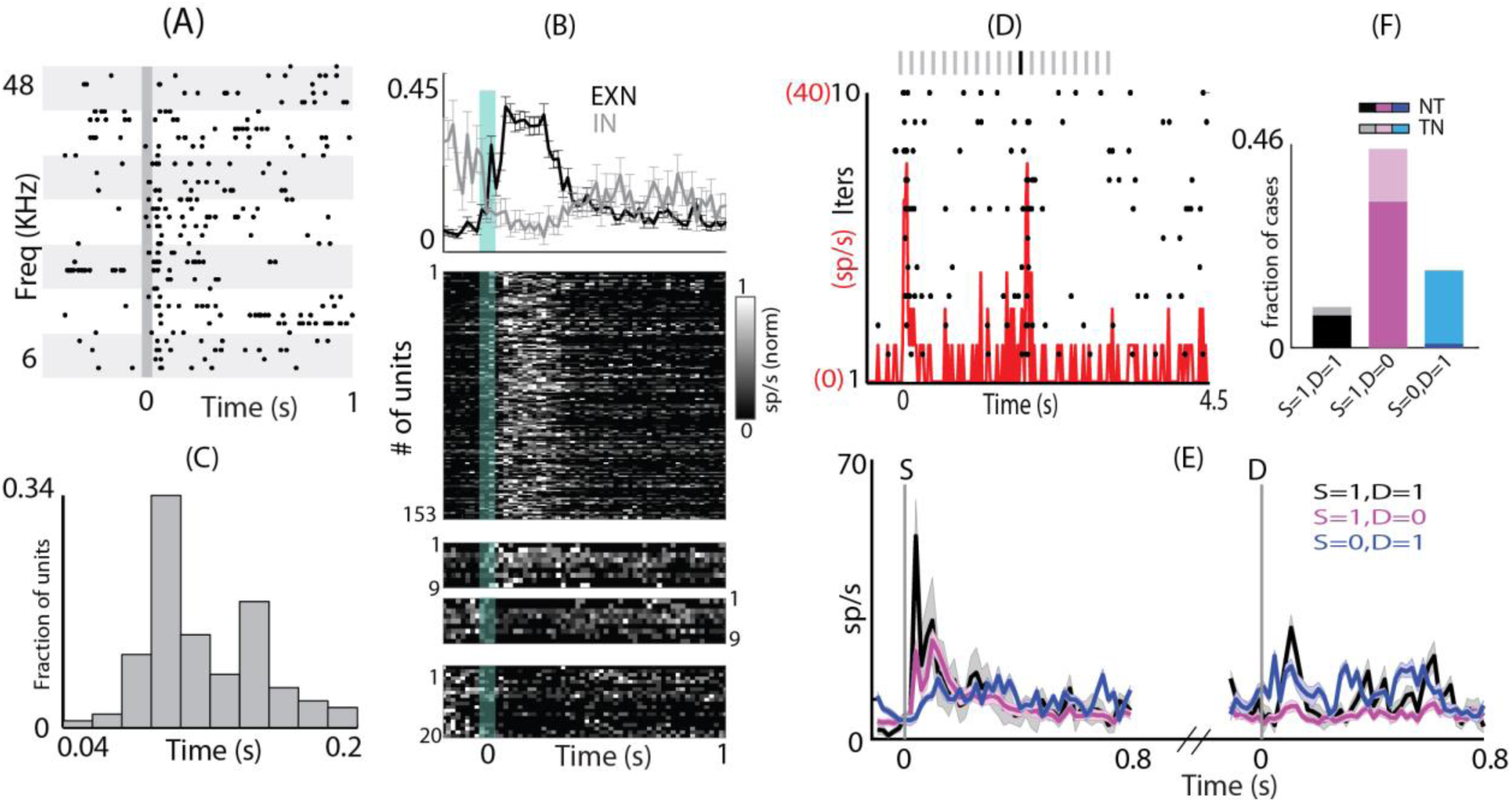
– Awake mouse OFC also shows deviance detection (A) Raster plot of an example unit in the OFC showing responses to tones of different frequencies in awake condition. (B) Top: mean normalized PSTH ± SEM of all units showing excitation (black) and inhibition (grey) upon auditory stimulation. Bottom: individual unit’s PSTH (upper panel: units showing excitation, middle two panels: units showing excitation to some frequency and inhibition to some other frequency, lower panel: units showing inhibition) (C) A distribution of peak response latency in the awake condition. (D) Raster plot (black) and PSTH (red) of an example unit showing responses to odd ball stimulus. (E) Mean population PSTH ± SEM of units responding to both standard onset and deviant (S=1, D=1; black), responding only to standard onset (S=1, D=0; magenta) and responding only to deviant (S=0, D=1; blue). (F) Fraction of units belonging to different categories as described in (E). The darker shades show the fraction of units belonging to NT and lighter shades show fraction of units belonging to TN group.

In the awake head-fixed passively listening mice, responses to TN and NT pairs of oddball stimuli were collected from 54 units (108 cases, Fig. 5D and E), which showed responses to at least one of the four stimuli (two S_1_ tokens and two D tokens), as earlier. The population PSTHs (Fig. 5E) clearly show that neurons responded only to the first token S_1_ and then to the D token and not to the other tokens, as was observed in the anesthetized OFC. Thus the basic observation of immediate adaptation to a single token and response only to deviant token, if at all, giving rise to deviant selectivity, is the same as in anesthetized mice. However, the responses in the oddball case were far more selective, sparse and context dependent in the case of passively listening mice than in anesthetized mice. In 30/108 (28%) cases there were responses to neither S_1_ nor D, which is significantly larger than that in the anesthetized OFC and AC (34/218, p<0.01 and 12/124, p<0.001 respectively, *two proportion z-test*). In 10/108 cases there were responses to both S_1_ and D (8 NT and 2 TN), 49/108 cases to only S_1_ (36 NT and 13 TN) and 19/108 cases to only the D token (1 NT and 18 TN, Fig. 5F). Thus overall, in the awake condition, there were responses to the D token in 27% (29/108) cases compared to 53% cases in the anesthetized OFC (Fig. 3B), showing even higher selectivity of responses. When considering the same sound tokens as S and D, unlike the anesthetized case we found far stronger context dependence of responses, particularly to tones. In 1(14), 14(30), 8(6) and 31(4) out of 54 units, there were responses to tones (noise), both as S_1_ and D, only as S_1_ and not as D, only as D and not as S_1_ and to neither S_1_ nor as D, respectively. Thus responses to tones were far sparser and selective in the oddball stimuli with only 23/54 units showing responses to the tone (either as S_1_ or as D), although for pure tones there were responses in 73% units (p<0.001, *two proportion z-test*).

### OFC receives the major projections from the dorsal AC but the major excitatory drive from the ventral AC

With our observed auditory response properties in the OFC single neurons, and its capability in modifying auditory cortical responses (Winkowski et al., 2013), it is important to decipher the sources of auditory inputs to the OFC to understand how such response properties are derived and how the circuit maybe involved during behavior. While the mouse OFC is being studied extensively in the context of behavior with different sensory stimuli and cues (Bissonette et al., 2008; Graybeal et al., 2011; Ward et al., 2015; Jennings et al., 2019; Liu et al., 2019), mouse OFC afferents that can carry auditory information is not clear. In fact, parallels of prefrontal structure and function between the mouse and the non-human primate appear to be inconsistent across studies, rodent atlases (Preuss, 1995; Laubach et al., 2018), although the mouse OFC (and PFC) receives medio-dorsal nucleus (MDN) inputs (Groenewegen, 1988; Murphy and Deutch, 2018). With the rise in use of mice, due to genetic and other technical advantages, in frontal cortical research, it becomes important to characterize the anatomical inputs. In our case, the recording location is the lateral and ventral OFC (LO/VO) relatively in the rostral part of the OFC extent (Fig. 1A). Although in the rat, an analogous region is known to receive auditory cortical projections (Murphy and Deutch, 2018), anatomical and functional details of the sources are not known in the rodent.

To find the main source of auditory afferents in the OFC that would be capable of driving auditory responses, we performed neuroanatomical experiments by injecting 200 nl green retrobeads (Lumafluor Inc) stereotactically in the OFC. We specifically targeted the location where we perform electrophysiological recordings (*n* = 9, Figs. 1A&6A). Number of beads in the regions across the rostro-caudal (RC) and medio-lateral (ML) extent encompassing the ventral AC (AuV), primary AC (A1) and dorsal AC (AuD) were observed and quantified (Fig. 6B, Methods). In a subset of experiments (Methods) we also confirmed the extent of A1 and AuV with m-cherry labeled projections (Fig. 6C) from the ventral division of the medial geniculate body (MGBv). Since both AuV and A1 (Fig. 6C) receive MGBv projections (Ohga et al., 2018), the region dorsal to A1 could be identified as AuD. The regions were demarcated in other mice (without MGBv labeled projections) based on the mouse atlas (Paxinos and Franklin, 2013), which corresponded well with our observations in mice with MGBv injections. Similarly, labeled MGBv projections also allowed us to corroborate lamina specific distribution of cells projecting to the OFC. The average projection profile across the ML extent of AC showed that AuD had the highest density of cells projecting to the OFC, followed by AuV, with minimal projections from A1 (Fig. 6D). The lamina specific distributions (Fig. 6E) showed that most of the projections to OFC from the AC originated in the infra-granular layers (Layer V/VI). Thus with both A1/AuV and AuD projecting to the OFC, it is likely that both the lemniscal and nonlemniscal pathways are involved in shaping auditory responses in the OFC.

**Figure 6.**
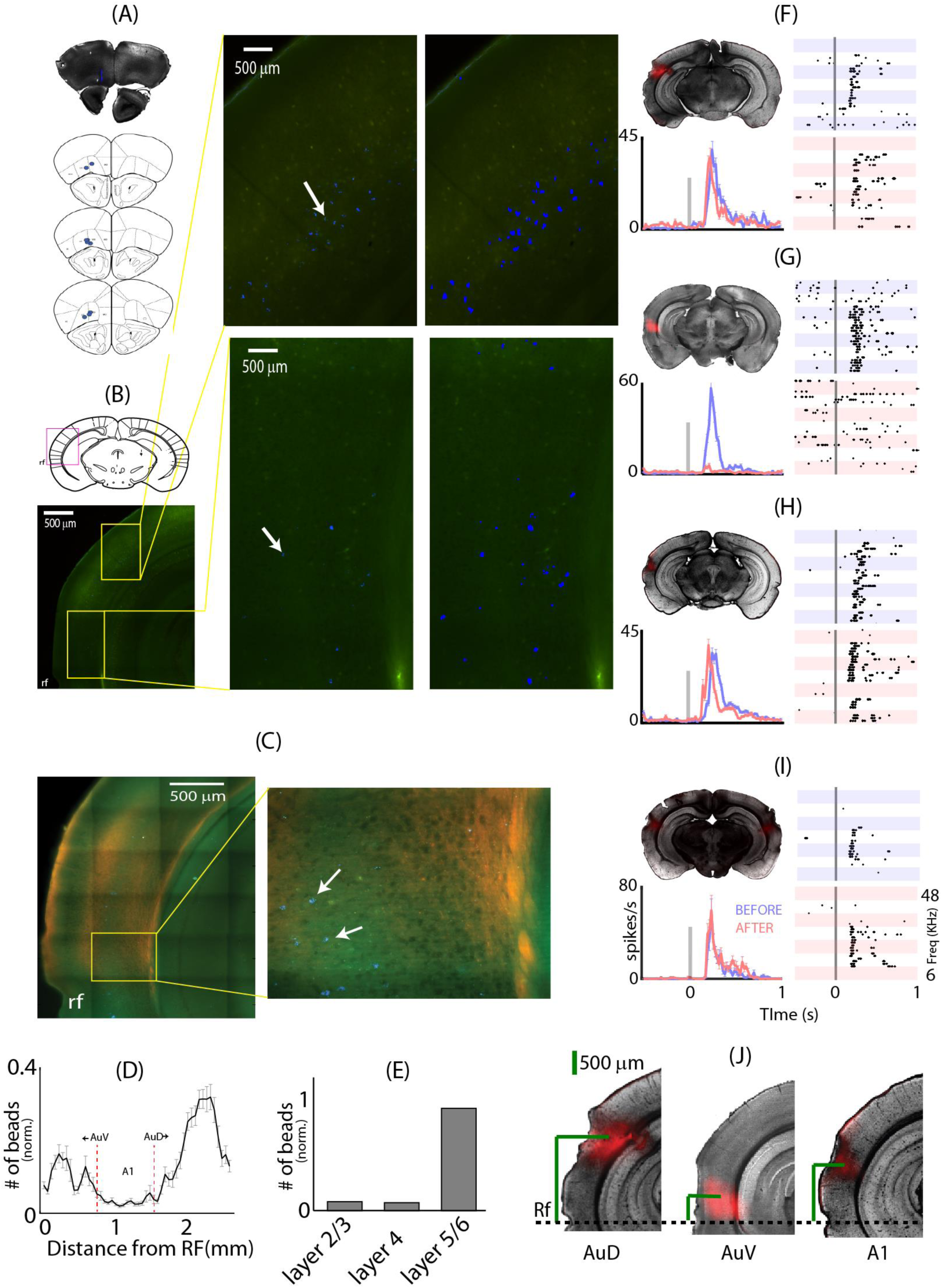
– Distinct contributions of AC divisions in OFC auditory responses (A) Top: Coronal brain section showing retrobeads injection site (white arrow) in the OFC. Bottom: Injection sites in all 9 animals (B) Top left: coronal brain section from mouse atlas showing auditory cortex. Magenta box roughly marks the part of the brain region shown in bottom left image. Bottom left: regions of AuD and AuV are marked in the yellow box. rf: rhinal fissure. Middle: AuD (top) and AuV (bottom) showing labeled cell bodies (white arrows) by the retrograde transported beads from OFC. Right: Same regions with enhanced intensities of the blue pixels for easy visualization of the beads. (C) Left: Brain section showing layer 4 of the AC labelled with AAV-mcherry injected in the MGBv. Right; zoomed in image of the area inside the yellow box on the left showing beads. (D) Mean number of beads as a function of distance from the rf. Dashed red lines mark the extent of A1 (E) Laminar distribution of beads in the AC (F) Brain section showing block site in the AuD; Raster plots of an example unit before and after silencing the AuD; mean population PSTH ± SEM before (blue) and after (red) silencing the AuD. Vertical grey line is the stimulus time. (G) Brain section showing block site in the AuV; Raster plots of an example unit before and after silencing the AuV; mean population PSTH ± SEM before (blue) and after (red) silencing the AuV. (H) Brain section showing block site in the A1; Raster plots of an example unit before and after silencing the A1; mean population PSTH ± SEM before (blue) and after (red) silencing the A1. (I) Brain section showing dual block sites in both A1; Raster plots of an example unit before and after silencing both the A1s; mean population PSTH ± SEM before (blue) and after (red) silencing both the A1s. (J) Block sites in the AuD, AuV and A1 from the rf.

We next tested the functionality of projections from AC to the OFC, to determine the contribution of the different regions of AC to the auditory responses in the OFC. In a series of experiments, single unit recordings with responses to pure tones in the OFC were performed before and after pharmacological inactivation of AuD (*n* = 3), AuV (*n* = 5) and A1 (*n* = 7) with muscimol and baclofen (Methods). The site of inactivation was confirmed by tracking fluorescent SR101 (Fig. 6F-H, left top) mixed with the muscimol-baclofen solution.

Contrary to expectation, inactivation of AuD (Fig. 6F), did not affect the single unit OFC responses to auditory stimulation (Fig. 6F, right, example single unit dot raster). The population mean PSTHs (*n* = 45 units) before and after AuD inactivation (Fig. 6F, bottom left), showed no difference (blue and red, mean spike rates 72.2+/-3.7 and 65.7+/-4.8, NS). All frequencies responding significantly before inactivation were included in constructing the population PSTHs. While the AuV, with lesser number of neurons than AuD projecting directly to the OFC (0.11+/-0.02 & 0.2+/-0.03 beads, p<0.01 *t-test*; normalized), was found to be the source of almost the entire auditory driven excitatory activity in the OFC. Similar plots as before, of example dot raster and population mean PSTHs (*n* = 43 units) for before and after AuV inactivation (Fig. 6G) show that OFC auditory responses were almost completely abolished following AuV block (98.7+/-3.3 and 14.9+/-1.4, p<10^-76^). Of course, other than the direct inputs to OFC from AuV, other inputs to OFC providing such excitatory auditory inputs cannot be ruled out, but such indirect pathways also must originate from the AuV. Thus the dorsal and ventral divisions of nonprimary AC are in stark contrast of each other in terms of their contribution to OFC auditory responses. However, their direct anatomical connections show characteristics opposite to their functional contributions. Similar to AuD block, inactivation of A1 (Fig. 6H) did not lead to any change in firing rates (70.1+/-2.4 and 72.9+/-3.4, NS), as observed in population mean PSTHs (*n* = 62 units) and dot raster plots (Fig. 6H). The possibility of a contralateral A1 contribution through indirect pathways to OFC was also considered. Bilateral inactivation of A1 (Fig. 6I, top left) also did not change firing rates of OFC neurons as assessed through population mean PSTHs (*n* = 30 units) and mean firing rates of single units in response to tones before and after pharmacological inactivation of both A1s (61.7+/-10.07 and 79.5+/-11.3, NS).

### Auditory responses in the OFC originate from both, the lemniscal and nonlemniscal medial geniculate body (MGB)

Although AuV, alternately A2 (Stiebler et al., 1997), is classically considered a belt area (Kaas and Hackett, 2000), the presence of lemniscal inputs to AuV (Fig. 6C, Ohga et al., 2018) opposes the classical view that MGBv projects exclusively to A1/AAF in the core AC. Since A1 and AuD inactivation did not cause changes in response rates in the OFC and given MGBv projections on AuV, we hypothesized that OFC auditory responses originate in the lemniscal MGBv. To test the hypothesis, single unit recordings in response to pure tones in the OFC were performed before and after MGBv inactivation (Fig. 7A). As with AuV inactivation, auditory responses in the OFC were completely abolished with MGBv block (23 units, 121.3+/-9.7 and 18.5+/-1.5, p<10^-16^). Since MGBv efferents almost entirely project to A1/AAF and AuV, we conclude that the entire auditory driven excitatory input originates in the lemniscal auditory thalamus (MGBv). For our MGBv inactivation we confirmed *post hoc* (Fig. 7A top left, inset) the lack of SR101 in dorsal MGB (MGBd) (Paxinos and Franklin, 2013) to rule out inactivation of the neurons projecting to AuV from there. Physical damage to MGBd during GABA agonist injections to MGBv was also ruled out by repeating the experiments with saline injections to MGBv which did not alter tone response rates in the OFC (*n* = 18 units, 93+/-6 and 75.6+/-6.6, NS). Also, since AuD with major thalamic afferents from MGBd (Kok and Lomber, 2017) did not alter OFC responses, involvement of the nonlemniscal MGBd in OFC auditory responses are at best minimal. The contribution to OFC auditory responses of the remaining major nucleus of the auditory thalamus (Calford, 1983; Rouiller et al., 1989; Winer et al., 1999), the medial MGB (MGBm), was also tested in a similar fashion (Fig. 7B). As opposed to abolition or no change in response rates observed in the other inactivation experiments, responses to pure tones altered dramatically in the OFC following inactivation of MGBm. Most units showed a behavior similar to the example single unit activity before (blue) and after (red) MGBm block shown in Fig. 7B (right). The population mean PSTHs (42 units) showed no change in peak response rates (54.7+/-2.8 and 52+/-2.8, NS, *t-test*) with MGBm inactivation, but the responses became persistent, with spiking continuing following the stimulus onset sometimes up to 1s and mean response duration almost doubling (177+/-6 and 314 +/-11 ms, p<0.001, Fig. 7C). Thus, MGBm, which has multiple sources of auditory inputs as well as other sensory inputs and crucially involved in auditory fear conditioning, in the non-lemniscal auditory pathway, is a source of long lasting auditory driven inhibition in the OFC. The net inhibitory effect in OFC originating from the MGBm thus sharpens auditory responses. Likewise, when the inhibitory source in MGBm is itself inhibited, based on behavioral relevance of stimuli (O’Connor et al., 1997) the effect would be to convert a short duration response into a long lasting persistent activity, a feature of prefrontal cortical responses in working memory (Fuster and Alexander, 1971; Funahashi et al., 1989; Schoenbaum and Setlow, 2001).

**Figure 7.**
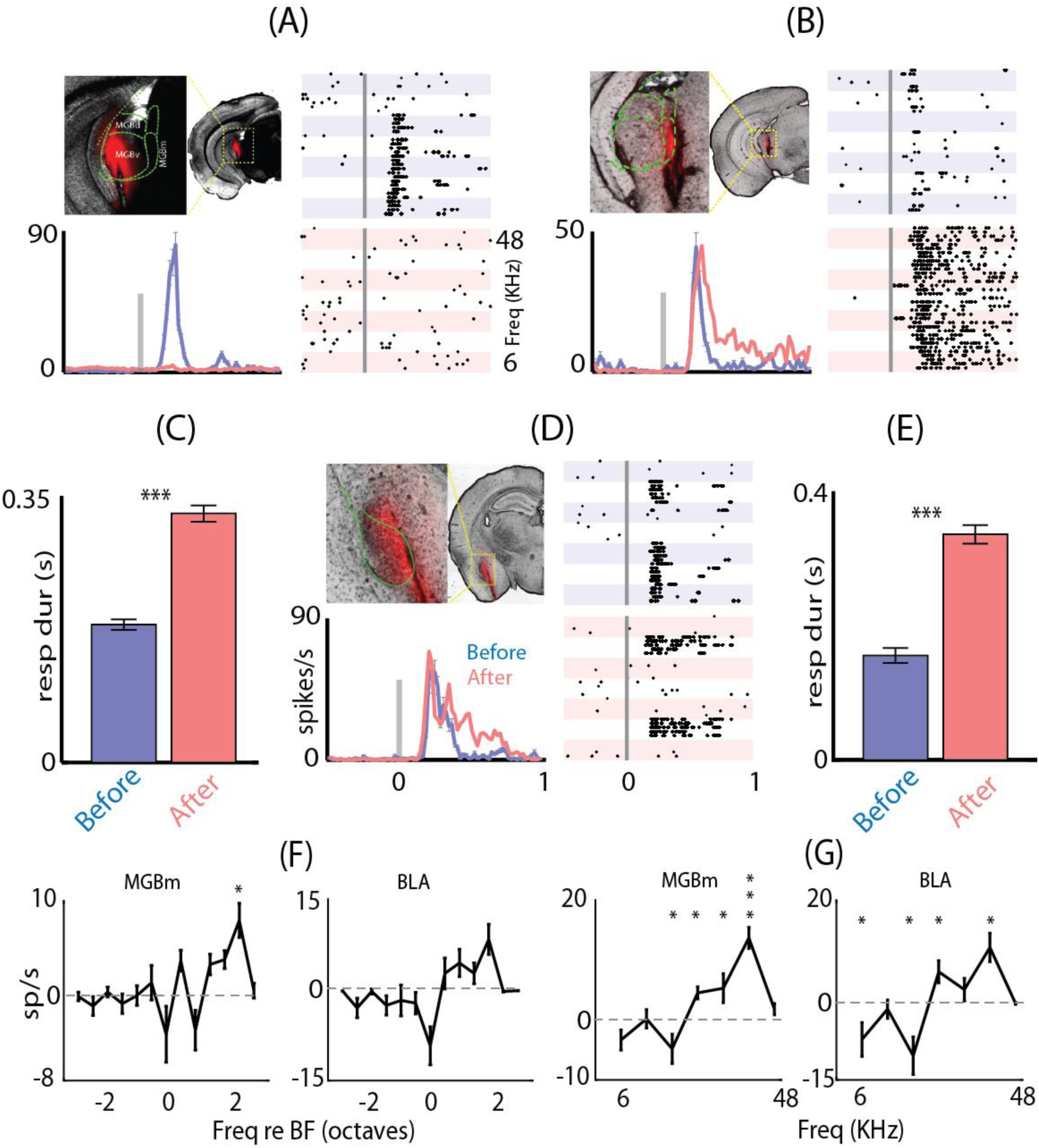
– Parallel excitatory and inhibitory contributions to OFC auditory responses originate in the lemniscal and non-lemniscal auditory thalamic nucleii (A) Brain section showing block site in the MGBv; Raster plots of an example unit before and after silencing the MGBv; mean population PSTH ± SEM before (blue) and after (red) silencing MGBv. Vertical grey line is the stimulus time. (B) Brain section showing block site in the MGBm; Raster plots of an example unit before and after silencing the MGBm; mean population PSTH ± SEM before (blue) and after (red) silencing the MGBm. (C) Mean response duration before and after silencing the MGBm (D) Brain section showing block site in the BLA; Raster plots of an example unit before and after silencing the BLA; mean population PSTH ± SEM before (blue) and after (red) silencing the BLA. (E) Mean response duration before and after silencing the BLA. Mean inhibitory inputs into OFC, from MGBm and BLA as a function of (F) frequency with respect to BF and (G) absolute frequency.

Further investigation of the nonlemniscal source (MGBm) of auditory responses in OFC, requires following up on its broad projections across entire AC, multimodal and limbic cortical areas (Lee and Winer, 2008; Lee, 2015) through cortical layers 1 and 6 (Huang and Winer, 2000) and the amygdala, specifically the lateral amygdala, LA (Iwata et al., 1986; LeDoux et al., 1990; Woodson et al., 2000). Given the strong connection from MGBm to LA (Woodson et al., 2000) and LA to BLA (Ledoux, 2000) and dense projections from BLA to OFC (Lichtenberg et al., 2017), we hypothesized that the major MGBm contribution to OFC auditory responses is through the BLA. Barring the possible auditory response contributions in the OFC of MGBm through AC layer 1 (Huang and Winer, 2000) and AC to LA through BLA, effects of BLA inactivation would produce an effect similar to that of MGBm silencing. As expected, on inactivating the BLA (Fig. 7D) we found a similar response pattern with long lasting persistent activity in the OFC (example dot raster, Fig. 7D, right), with no change in peak firing rates as assessed through the population mean PSTHs before and after BLA block (56 units, 68.7+/-4.6 and 78.9+/-5, NS, *t-test*). As with MGBm block response duration increased (Fig. 7E) by similar degrees with BLA inactivation (response duration: 156+/-11 and 338+/-14, p<0.001, *t-test*).

The changed response to tones after MGBm or BLA inactivation was quantified by considering the difference in mean rate responses (after-before) to the different frequencies (in the window of response duration after inactivation, Methods) relative to the BF of the units (Fig. 7F, MGBm, block left, BLA block, right). In both cases similar pattern of rate difference was observed, with almost all frequency components relative to BF, showing no significant difference except a peak at 2 octaves above BF and a negative peak at BF. Thus the MGBm or BLA based inhibition into the OFC is not organized in a BF specific way. Rather when considering the changes in firing rate before and after MGBm or BLA inactivation (Fig. 7G, left and right) in absolute frequency it is found that significant inhibitory inputs are in the middle frequency region of mouse hearing (17-34 kHz). Further we can conclude that there is a trend of a significant net excitatory contribution mediated through both MGBm and BLA in the lower frequencies (6-12 kHz). Since the pathway originating in the MGBm is associated with fear conditioning it is likely that this pathway is already shaped based on the natural previous experience of fear associated stimuli and auditory events.

### Auditory response properties and deviant selectivity of OFC: contributions of auditory input sources

Our inactivation experiments of the different divisions of the AC, the MGB and the BLA conclusively show the sources of the major auditory driven effective excitatory and inhibitory inputs to the OFC. The conclusions are based on the overall firing rates and duration of responses to tones. Further analyses of the responses before and after inactivation of A1, AuD, MGBm and BLA (the cases in which responses were not abolished) show how detailed temporal aspects of OFC responses, important in population coding based on synchronization and ultimately in contributing to plasticity and adaptive coding, are derived from the different sources. Since AuD inactivation did not cause any changes in responses in the OFC, changes in other aspects of responses were not considered further.

We find that, with inactivation of A1, although there are no changes in firing rate, there is a significant reduction in latency of responses to tones in single units of the OFC (306+/-3.4 ms, 258+/-4.33 ms, p<0.001, *paired ttest*, Fig. 8A, left). Such latency reduction is usually associated with, a stimulus getting effectively stronger (for example with increasing sound level of noise, Fig. 1B), weaker long term adaptation (Fig. 2E) or a disinhibition. Similarly, on considering latencies before and after inactivation of MGBm and BLA, opposing effects of latency were found, although in both cases similar changes were observed in terms of response rates and response duration. MGBm inactivation led to marked increase in latency to tones (246.5+/-4.4, 296.6+/-7.3 ms, p<.001, *paired ttest*, Fig. 8A, middle) while inactivation of BLA barely led to a reduction in latency (315.4+/-7.4, 297.4+/-8 ms, NS *t-test*, removing effect of outliers and comparing medians, p<0.05 *ranksum*). Since the initial latency of response varied between the two populations of single units (before MGBm block and before BLA block) we also compared the fractional change in latency of single units on MGBm inactivation and BLA inactivation (mean 27% and 7 % *unpaired t-test* p<0.001). Thus the effective inhibition in the OFC, originating from MGBm is not simply relayed by the BLA. It is likely that there are other auditory sources that also provide inhibition via BLA on to OFC. Such sources of inhibition on OFC through BLA, likely from AC, (Bertero et al., 2019) remain intact on MGBm inactivation leading to longer latency, while with BLA inactivation their effect is removed to lead to the reduced latency. A likely candidate pathway would be A1/AuV to the lateral amygdala (LA, Romanski and LeDoux, 1993; Romanski et al., 1993; Tsukano et al., 2019), which in turn projects to the BLA (Janak and Tye, 2015). Further, LA also receives inputs from MGBm (Ledoux, 2000; Woodson et al., 2000) and is involved in fear conditioning, and thus it is a critical pathway to provide affective value information associated with particular auditory stimuli to the OFC.

**Figure 8.**
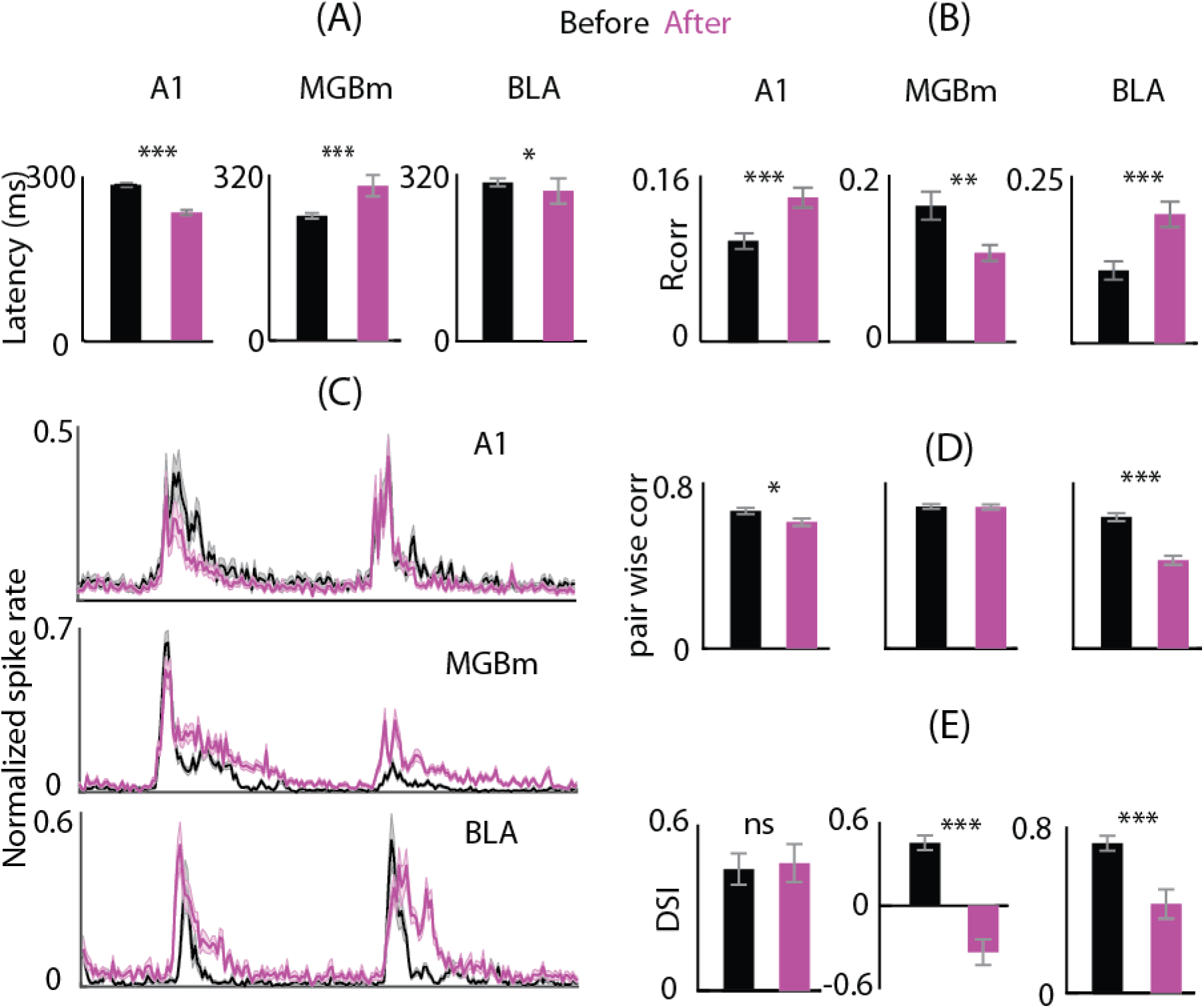
– Both OFC deviant selectivity and spike time based response properties are shaped by the nonlemniscal pathway unlike by A1 (A) Mean population latency ± sem to pure tones before (black) and after (magenta) silencing A1 (left), MGBm (middle) and BLA (right). (B) Mean population reliability (R_corr_) ± SEM before (black) and after (magenta) silencing A1 (left), MGBm (middle) and BLA (right). (C) Mean population PSTH ± SEM in response to odd ball stimulus before (black) and after (magenta) silencing A1 (top), MGBm (middle) and BLA (bottom). (D) Mean population pairwise correlations ± SEM before (black) and after (magenta) silencing A1 (left), MGBm (middle) and BLA (right). (E) Mean population DSI ± SEM before (black) and after (magenta) silencing A1 (left), MGBm (middle) and BLA (right).

Spike timing is crucial in generating synchronization in populations of neurons (Kreuz et al., 2007; Ermentrout et al., 2008), plasticity (Benedetti et al., 2009; Li et al., 2014) and in forming associations (Atilgan et al., 2018). Such temporal aspects of auditory responses are important in adaptively modifying and transforming sensory stimulus driven responses to behavior associated responses in the OFC. We first considered spike timing reliability in the OFC single units before and after inactivation of the different auditory sources. A1 inactivation remarkably increased spike timing reliability quantified through R_corr_ (Methods, from 0.0953±0.01 to 0.1385±0.01, p<0.001, *paired t-test*, Fig. 8B, left) in response to tones. Again, a differential effect was observed with BLA and MGBm inactivation. While MGBm inactivation decreased spike timing reliability (from 0.16+/-.02, to 0.11+/-0.01, p<0.01, *paired t-test*, Fig. 8B, middle), BLA inactivation nearly doubled reliability of spike times (from 0.11+/-.02, 0.19+/-.02, p<0.001, *paired t-test*, Fig. 8B, right) in repeated presentation of tones. Since inhibition is crucial in precise spike timing (Pouille and Scanziani, 2001; Wehr and Zador, 2003), and with MGBm having the highest reliability in spiking (Anderson and Linden, 2011) among the auditory thalamic nuclei, the loss of BLA mediated feedforward inhibition originating from MGBm, leads to the decreased reliability is spiking in the OFC. However, since A1 acts as a source of spike timing jitter in the OFC (Fig. 8B, left), blocking of BLA also effectively removes the effect of A1 dominated spiking jitter introduced in the OFC through LA.

To analyze the contribution of the different auditory sources to response properties of populations of OFC single units. We considered the case of NT and TN oddball stimuli since the hallmark of the auditory responses of the OFC is context dependence and deviant detection (Fig. 8C). Pairwise correlations (Methods) can strengthen coding efficiency (Hung et al., 2015) and enhance plasticity (Feldman, 2009). Thus, synchronization of responses in pairs of simultaneously recorded OFC single units was considered before and after inactivation of each of the auditory sources. Inactivation of A1 produced a small decrease in synchronization with oddball stimuli (from 0.67+/-0.01 to 0.61+/-0.02, p<0.05, *t-test* Fig. 8D, left). However, MGBm inactivation caused no change in pairwise correlations (0.69 +/-0.01 and 0.69+/-0.01, NS, Fig. 8D, middle) while BLA inactivation reduced pairwise correlations (from 0.64+/-0.02 to 0.43+/-0.02, p<0.001 *t-test*, Fig. 8D, right). Thus the effective inhibitory input to the OFC from the BLA synchronizes activity across populations of single units, independent of the MGBm input to the BLA. As BLA inactivation increases spike timing reliability of single neurons, it suggests that auditory driven indirect AC input to the BLA on to OFC, causes synchronization across pairs of neurons by reducing across trial similarity in responses of single neurons, enhancing a population level representation. The above is further corroborated by the fact that A1 inactivation reduced pairwise correlations in the OFC, although to a lesser degree. Finally, we consider the contribution of different auditory sources on the deviant detection property of OFC single units. Since we did not have sufficient data on pairs of NT and TN both before and after inactivation of different structures, we considered DSI (Methods) as opposed to CSI to quantify oddball selectivity. Population mean PSTHs of OFC single units in response to oddball stimuli before and after inactivation of A1, MGBm and BLA (each row, Fig. 8C) show emergence of persistent activity at both the onset and the deviant except in the case of A1 block. Quantification of the selectivity (DSI) before and after block shows that A1 does not contribute to DSI in the OFC, while inactivation of both the MGBm and the BLA produced drastic reduction in DSI (from 0.44+/-0.06 to 0.46+/-0.07, p=0.57, 0.46+/-0.05 to −0.34+/-0.09, p<0.001 and 0.72+/-0.04 to 0.43+/-0.07, p<0.001 respectively *t-test*, Fig. 8E). The difference in reduction of DSI in case of MGBm and BLA inactivation, suggest that the primary source of the selectivity to oddball stimuli in the OFC is the MGBm, while it is further strengthened by auditory inputs from the BLA from sources other than MGBm (like AC) to the OFC. However, it should be noted that, with neither MGBm nor BLA inactivation responses to subsequent tokens after the onset token appeared. So although values of DSI reduce on average to produce lower oddball selectivity in both cases (MGBm and BLA inactivation), the OFC’s intrinsic selectivity to deviant (onset token also a deviant following silence) and subsequent fast adaptation over a long time scale is unchanged. AuV is the source of the primary excitatory input to the OFC and the AuV itself does not show deviant selectivity (Fig. 3C). Thus adaptation in synapses providing the auditory input and the recurrent synapses within the OFC along with the membrane properties of OFC neurons themselves likely give rise to the pure oddball detection responses in the OFC.

## Discussion

We find the presence of robust auditory responses in single units of the mouse OFC during passive listening both in awake as well as in the anesthetized state. Auditory sensory responses in the OFC as in AC showed robust responses to tones and noise, the usual characterization stimuli. Other studies probing the OFC with different kinds of auditory stimuli in the mouse (Winkowski et al., 2017) and in primates (Rolls et al., 2006) have also found robust responses. However, in our study we show that the OFC auditory responses were drastically different from AC responses in a number of major ways, which have not been studied in the OFC previously. Although AC auditory responses are also affected by multiple time scales of adaptation (Ulanovsky et al., 2004), OFC neurons showed extremely long history dependence and adaptation lasting above 10s as observed with comparisons of OFC and A1/AuV responses with variety of ITIs (Fig.2). This difference was visible when ITI was varied between tone presentations and shorter ITIs produced higher latencies and lower peak spike rates as compared to longer ITIs in the OFC. The effects of such adaptation were stronger in the awake as there were far fewer responses to deviants in TN and NT oddball stimuli. The difference in history dependence between OFC and AC thus potentially encodes the sensory aspect of the auditory memory over long timescales. Encoding auditory memory is essential in the OFC as it is known to play a key role in creating various stimulus-outcome associations (Delamater, 2007; Rudebeck et al., 2008; Sadacca et al., 2018) and their revaluation. Since the outcome of a stimulus is almost always temporally offset by large durations (Pavlov, 1927; Fuster and Alexander, 1971) such long history dependence is essential in creating associations between the two as in auditory trace conditioning (Runyan et al., 2004; Gilmartin and McEchron, 2005). The OFC neurons show pure deviance detection during oddball stimulus streams, not responding to any repetition of the standard stimulus except the first one. SSA is present in the auditory pathway as early as the inferior colliculus, MGB as well as AC (Antunes et al., 2010; Taaseh et al., 2011; Ayala and Malmierca, 2013) where there is a gradual reduction in the spike rates to the repeating standards due to adaptation but here in the case of OFC we find a complete and sudden cessation of responses right from the first instant of repetition such that CSI distribution when computed with S_ALL_ and S_P_ were not significantly different unlike seen in the AC (Fig. 4). This faster and stronger adaptation to repetition appears to emerge in the OFC possibly achieved by local circuits within OFC through recurrence (Yarden and Nelken, 2017). The CSI distribution when computed with S_1_ shows a negative mean showing that responses in the OFC are highest to the biggest change in the stimulus space as the first token of the standards in the sequence to be the biggest change.

The presence of long timescales of adaptation in an anesthetized preparation shows the intrinsic nature of the OFC neurons. In the awake mouse such temporal dependence is more pronounced with fewer responses to the oddball stimulus (Fig. 5F). Occurrence of a stimulus from silence (also a deviant) is sufficient in many cases to suppress responses to a changed stimulus (oddball) within 2 seconds of the first stimulus token (Fig. 5E&F). However, as evident in our tuning data in the awake mice, where single tokens of different tones are presented with a gap of 5s, single units in the OFC respond to a changed stimulus (Fig. 5A). Thus in the awake condition stimulus changes beyond 2 seconds and within 5 seconds are detectable, thus making the history dependence more pronounced than in the anesthetized state.

The long time scale adaptation or history dependence observed in the OFC has also been observed to a similar degree in the anesthetized rat MGBm (Antunes et al., 2010), where SSA is observed in some neurons with up to 2s stimulus onset asynchrony (SOA). However, as shown by MGBm inactivation, it serves as a source of a long lasting inhibitory drive to the OFC. Thus, the long time scale dependence observed in the OFC, with suppression of responses beyond the onset stimulus cannot be derived by the same pathway from the MGBm, as the OFC’s main excitatory auditory drive’s source is AuV (Fig. 6G) from MGBv (Fig. 7A), which does not show such remarkable history dependence.

In our pharmacological block experiments we found that, while OFC robustly responded to auditory stimulation, A1 does not contribute to OFC response strength but only induces more jitter in spike timing and longer response latency possibly via an early inhibition through a weak input along the A1/AuV-LA-BLA (Romanski et al., 1993; Ledoux, 2000; Tsukano et al., 2019) or directly from A1 to OFC pathways (Fig. 6D & 6H). Anatomically, we found that secondary areas of the AC send most of the projections to OFC within AC with AuD showing the strongest labelling. Despite the strongest labelling, the OFC responses were not affected upon inactivating AuD. The mouse AuD is more involved in representing perceptual meaning of primarily temporally structured sounds while AuV is thought to represent value in terms of novelty (Weible et al., 2014; Geissler et al., 2016). The OFC is involved in stimulus outcome value computation, which is consistent with the result that AuV drives excitatory auditory responses in the OFC with basic tone and noise stimuli in the anesthetized state. AuD on the other hand is likely recruited to provide inputs to the OFC in a more behavioral context specific manner with complex (vocalizations) or multimodal stimuli (Morrill and Hasenstaub, 2018). The function and necessity of the AuD projections on to the OFC requires further investigation. Since OFC responses were abolished on silencing MGBv as with inactivation of AuV which receives direct inputs from the MGBv (Fig. 7A) (Ohga et al., 2018), we hypothesize that the OFC auditory responses are driven by MGBv via AuV.

Thus, we find that the auditory inputs to the OFC originate in at least two parallel regions in the MGB, the ventral and medial divisions. These two streams converge in at least two locations, the amygdala and the OFC. The input from AuV can be hypothesized to carry in sensory information with context dependence which is sharpened through the long lasting deviant selective inhibitory drive originating from the MGBm and modified in the BLA and also through local recurrent connections in the OFC. Thus the MGBm and BLA both provide a saliency filtering of the sensory inputs to the OFC and suppress responses in OFC units following deviant/salient auditory events. Moreover, control of thalamic deviant detection response is modulated by attention through the thalamic reticular nucleus (TRN) which can further modify deviant detection in the OFC (Yu et al., 2009) by sharpening selectivity.

The pathway originating in the MGBm also provides a means of controlling the sensory driven activity by causing persistent activity in the OFC, if needed, allowing the sensory stimulus to be associated with other outcome related delayed signals related to reward (Thorpe et al., 1983; Hikosaka and Watanabe, 2000; Stalnaker et al., 2014), prediction error (O’Neill and Schultz, 2013) or punishment (O’Doherty et al., 2001; Windmann et al., 2006) required for reinforcement in acquisition as well as in reversal learning. The non-lemniscal MGBm or BLA inactivation causing the OFC responses to persist for a long duration (up to ∼1 sec), could be further longer in the awake state as we see lack of responses to even the deviant stimulus in the majority of cases. Such longer persistent activity is common in the PFC in the awake state in the context of behavior involving working memory (Fuster and Alexander, 1971; Funahashi et al., 1989; Curtis and D’Esposito, 2003) and also in the OFC conveying incentive value of cues (Gallagher et al., 1999; Tremblay and Schultz, 1999). However, mechanisms of origins of such persistent activity are unknown. Thus we hypothesize that the auditory input pathway originating in the non-lemniscal MGB, provides the behavior related control on the sensory stimulus through two ways - when active it provides sharpening of deviant detection in the OFC and when inactive it converts the short stimulus response into persistent activity. In the latter case, the final input to OFC along this pathway from BLA, causes decorrelation of response in pairs of OFC units and precise spike timing across trials, other than causing persistence and lowering of deviant detection. Precisely timed spiking in the persistent OFC activity would aid in plasticity (Markram et al., 1997; Bi and Poo, 1998; Dan and Poo, 2004; Feldman, 2009) required for associating the auditory stimulus with delayed outcome related signals. Similarly, decorrelation of activity, removing redundancies in the population, allows more possibilities of creating associations with the OFC stimulus driven activity and outcome signals. Frontal cortex is also known to code reward value and other cognitive aspects in spike times (Smith et al., 2019) and hence the decorrelation and precise spike timing would aid in forming associations.

The above detailed inhibitory control can be achieved by suppression of an auditory stimulus driven responses in the MGBm or BLA. MGBm has been shown to be the first station in the auditory pathway that shows changes in the firing rate due to task related stimulus associations (Birt et al., 1979) and forms part of the thalamo-amygdaloid component in the auditory pathway (Ledoux et al., 1986) and crucial in fear learning. A study showed presence of a cell type in the MGBm that silences itself during conditioned stimulus presentation (O’Connor et al., 1997). MGBm has been shown to initiate a feed forward inhibition in the LA (Woodson et al., 2000), which through BLA to OFC can cause persistent activity. The MGBm connections with the limbic-related nuclei in the amygdala can uniquely alter OFC response allowing affective and emotional aspects of aversive stimuli to be associated with outcomes or changed outcome signals (LeDoux et al., 1985, 1991; Iwata et al., 1986; Cruikshank et al., 1992). The LA region receiving MGBm inputs also receive convergent inputs from the A1/AuV (Romanski et al., 1993; Tsukano et al., 2019), which may allow synchronization and coalescing sensory responses to salient affective stimuli (Winer, 2006; Weinberger, 2011). Thus the MGBm driven control of OFC activity is likely to do with associations of fear eliciting stimuli (Weinberger et al., 1995; Weinberger, 2011). It is also reflected in the fact that the frequency profile of the inhibitory inputs from the MGBm via the LA-BLA-OFC pathway is innately tuned to the 30 kHz region (Fig. 7) which is also the frequency range of fear induced vocalizations.

Similarly the BLA, known to encode valence (Paton et al., 2006; Janak and Tye, 2015; Zhang and Li, 2018), to OFC connections have been shown to be important in a number of ways in disruption of decision making and reversal learning (Zeeb and Winstanley, 2013; Orsini et al., 2015; Lichtenberg et al., 2017; Groman et al., 2019). Particularly BLA projections to the OFC are known to enable stimulus cue triggered reward expectations and ablation of BLA neurons projecting to the OFC impairs reversal learning due to inability to use positive outcomes guided choices following reversal. We hypothesize that the underlying deviant selectivity of auditory inputs imposed by the amygdala and lack of precise disinhibitory control to generate persistent activity to associate with the newly rewarding or positive outcomes in the OFC after contingency reversal leads to observed dysfunctions stated above. Thus we propose that the MGBm via LA-BLA and BLA itself act as controllers of persistent activity required for stimulus and delayed outcome associations during reversal learning, for aversive and rewarding cues respectively.

## Acknowledgements

HKS thanks IIT Kharagpur, MHRD for Institute Fellowship, SB thanks Wellcome Trust DBT India Alliance for Grant No. IA/I/11/2500270, MHRD SSLS scheme, SRIC Challenge Grant and IIT Kharagpur for funding.

